# CeMbio - The *C. elegans* microbiome resource

**DOI:** 10.1101/2020.04.22.055426

**Authors:** Philipp Dirksen, Adrien Assié, Johannes Zimmermann, Fan Zhang, Adina-Malin Tietje, Sarah Arnaud Marsh, Marie-Anne Félix, Michael Shapira, Christoph Kaleta, Hinrich Schulenburg, Buck S. Samuel

## Abstract

The study of microbiomes by sequencing has revealed a plethora of correlations between microbial community composition and various life-history characteristics of the corresponding host species. However, inferring causation from correlation is often hampered by the sheer compositional complexity of microbiomes, even in simple organisms. Synthetic communities offer an effective approach to infer cause-effect relationships in host-microbiome systems. Yet the available communities suffer from several drawbacks, such as artificial (thus non-natural) choice of microbes, microbe-host mismatch (e.g. human microbes in gnotobiotic mice), or hosts lacking genetic tractability. Here we introduce CeMbio, a simplified natural *Caenorhabditis elegans* microbiota derived from our previous meta-analysis of the natural microbiome of this nematode. The CeMbio resource is amenable to all strengths of the *C. elegans* model system, strains included are readily culturable, they all colonize the worm gut individually, and comprise a robust community that distinctly affects nematode life-history. Several tools have additionally been developed for the CeMbio strains, including diagnostic PCR primers, completely sequenced genomes, and metabolic network models. With CeMbio, we provide a versatile resource and toolbox for the in-depth dissection of naturally relevant host-microbiome interactions in *C. elegans*.

**Dataset accession numbers:** Whole genome sequencing data (PRJNA624308); microbiome sequencing [PRJEB37101 and PRJEB37035]; data supplement on the GSA Figshare Portal.

## INTRODUCTION

While there is little debate that microbiomes exert broad influence on their hosts (McFall-Ngai 2014; Gilbert *et al*. 2018), less is known about the mediators of this influence. Often the complexity of the systems renders interrogation impossible. Model hosts address variation by controlling much of the environmental, genetic and dietary drivers of host-microbiome interactions (Fraune and Bosch 2010; Douglas 2019), but often overlooking the importance or extent of genetic and functional variation on the part of the microbiome. The greatest advances in understanding have emerged largely from binary tests of one host and one microbe under gnotobiotic conditions (Fischbach 2018). While certainly valuable, these types of experiments likely also oversimplify the system in a manner that limits ability to identify properties that emerge from collaborations and competitions between microbiome members and their natural host. Thus, there is a need to develop well-characterized, tractable systems that faithfully capture the complexity of these interactions and identity of the molecular drivers of microbiome impact.

To this end, *C. elegans* has emerged as a powerful high-throughput system for studying host-microbiome interactions (Zhang *et al*. 2017). This free-living nematode has many inherent strengths including a short life cycle of 3 days and lifespans of 3 weeks, a well-defined and transparent body plan, widely available resources and facile methods for forward and reverse genetics, plus a wealth of understanding of its biology and physiology (Girard *et al*. 2007; Frézal and Félix 2015). In the wild, *C. elegans* harbors a characteristic gut microbiome community that is recruited from its surrounding environment (Dirksen *et al*. 2016; Samuel *et al*. 2016; Berg *et al*. 2016b). Meta-analyses of these natural microbiomes highlight core membership of over a dozen bacterial families, including Gammaproteobacteria (*Enterobacteriaceae, Pseudomonadaceae*, and *Xanthomonodaceae*) and Bacteroidetes (*Sphingobacteriaceae, Weeksellaceae, Flavobacteriaceae*) (Zhang *et al*. 2017).

Here we establish a publicly available and well-defined model microbiome for use in *C. elegans* (CeMbio). This set is composed of 12 bacteria from 9 different families that represent the core microbiome of *C. elegans* based on analyses and empirical studies of intestinal colonization. These bacterial strains are presented with fully sequenced and annotated genomes, metabolic network reconstructions, and robust protocols for their use in *C. elegans* studies and beyond. All of the bacteria effectively colonize the *C. elegans* gut both alone and as a community, which can impact on the growth and development of the host. The CeMbio community has broad application to any aspect of *C. elegans* biology from aging to pathogenesis, development to neurobiology, and any aspect of physiology where a more natural environment is desired. Ultimately, pairing of this well-defined microbiome and highly-tractable host is envisioned to complement other systems (e.g., (Fraune and Bosch 2010; Brugiroux *et al*. 2016; Douglas 2019)) in advancing understanding of the mechanisms of microbiome impact on host health and disease.

## METHODS

### Bacterial collections of natural Caenorhabditis populations

The CeMBio strains were chosen as described in the next section from a set of previously cultured bacteria from the Félix, Samuel, Schulenburg and Shapira labs (Dirksen *et al*. 2016; Samuel *et al*. 2016; Berg *et al*. 2016a), plus an additional collection of ca. 140 bacterial strains also isolated from wild *Caenorhabditis* animals in the Félix lab (JUb130-274; Table S1).

For the new Félix lab collection, *Caenorhabditis* animals were collected from rotting fruit and stems from in and around Paris as well as Brittany and Indre (France). Substrate samples were brought back to the laboratory to isolate worms using adapted methods as in Barrière and Félix (2006). Briefly, while working aseptically, samples were plated onto sterile petri plates containing Normal Growth Medium (NGM: Autoclave 3g NaCl, 2.5g Bacto-Peptone, 17g Bactor Agar, 1L sterile water; after cooled to 55°C add 1 ml of 5 mg/ml Cholesterol, 1 ml 1 M CaCl_2_, 1 ml 1M MgSO_4_, 25 ml 1M pH6 KPO_4_) and diacetyl, a chemical attractant (10 ml of 1:30 dilution onto the agar at the opposite of the 90 mm plate). Nematodes were identified to the genus level immediately by morphology, and to the species level through subsequent crosses and molecular verification as needed (most common around Paris: *C. elegans, C. briggsae*, or *C. remanei*).

Animals were then surface sterilized following a method similar to that described in Portal-Celhay and Blaser (2012). Worms were washed off plates using sterile M9, then spun down for 2 minutes at 3,000 rpm. We removed excess liquid, then transferred worms to 55mm plates with 100 mM Gentamicin in NGM agar. After an hour on these plates, worms were washed off the plate with M9 and spun down for 2 minutes at 3,000 rpm. Excess liquid was pipetted off and the wash was repeated.

Gut-associated microbes were isolated from the surface sterilized samples above (JUb130-JUb265) or previously frozen nematode strains (JUb266-JUb274) using standard microbiological isolation techniques. We placed three adult worms from each sample into an Eppendorf tube with 500µl sterile water, then deadbeat (Mini-beadbeater, BioSpec Products, Bartlesville, OK, USA) them at maximum speed for two minutes. 100 µl lysed material from each sample was then plated onto each of four types of agar media in 90 mm plates. The liquid was spread and the plates were allowed to dry completely before wrapping them with parafilm. The four types of bacterial culture media included: NGM, Yeast Malt Extract Agar (YMEA), lysogeny broth supplemented with mannitol (LB+M: 5 g NaCl, 10 g Tryptone, 5 g Yeast Extract, 15 g Bacto-agar, 10 g Mannitol, 975 ml sterile water, 50µl 10N NaOH), and chitin agars (Autoclave: 20 g Agar, 4 g Chitin, 0.75g K_2_HPO_4_, 0.5g MgSO_4_ x 7H_2_O, 0.35g KH_2_PO_4_, 0.01 g FeSO_4_ x 7H_2_O, 0.001 g MnCl_2_ x 4H_2_O, 0.001 g ZnSO_4_ x 7H_2_O and 1 L sterile water). All media were supplemented with antifungals (20 ml/l nystatin and 0.05 g/l cycloheximide) after autoclaving and cooling to 55°C. Bacteria were cultured at room-temperature (23°C). Colonies were picked 1 to 3 days after bead beating and again 1 to 5 weeks after in an effort to isolate both slow and fast growing bacterial strains. Single colonies were picked again a few days later. Once they were in pure culture, bacterial strains were preserved in 10% glycerol solution and frozen at −80°C for long-term storage. Strains were identified by sequencing the 16S rRNA sequence using the following primers: 27f-1492r (Lane 1991), 530f-1391r (Walker and Pace 2007), S-C-Act_235a/878 (Stach *et al*. 2003), Act283f/1360r (McVeigh *et al*. 1996) (Table S1).

### Characterization and maintenance of CeMbio bacteria

Twelve bacterial isolates were selected as part of the CeMbio resource (Table 1). Maximum-likelihood phylogenetic analysis was used to inform candidate selection for the CeMbio resource by comparing a total of 510 sequences from *C. elegans*-related bacterial isolates from the Félix, Samuel, Schulenburg, and Shapira labs to the 12 most common OTUs, which we inferred by repeating our previous meta-analysis (Zhang *et al*. 2017) with only the natural worm samples (Figure 1A; Table S2). Taxonomic identity of the CeMbio strains was inferred via comparisons of genome-derived 16S rRNA sequences with same-family bacterial type strain sequences from the SILVA ribosomal RNA database project (Quast et al., 2013) as of 2018-11-20 using maximum-likelihood phylogenetic analysis (File S1). In both cases, we constructed multiple sequence alignments of the 16S sequences using the R package *DECIPHER* (Wright, 2016) and the aligner *SINA* (Pruesse et al., 2012). Phylogenetic tree reconstruction was performed with *IQ-TREE* (Nguyen et al., 2015) with its implementation of *ModelFinder* (Kalyaanamoorthy et al., 2017) for maximum-likelihood model selection and a total of 10000 ultra fast bootstrap (Hoang et al., 2018) replicates. The resulting trees were visualized with the R packages *ape* (Paradis and Schliep, 2019), *ggplot* (Wickham, 2009), and *ggtree* (Yu et al., 2017). An isolate was assigned to a particular taxon if it clustered in a clade containing only strains of a single species or genus with at least 75% bootstrap support (10,000 replicates). Using this approach, we could assign all strains to a genus and seven of them to a species (Table 1). Using a complementary phylogenomics approach (see genomics section below) a total of eight strains could be given strains designation and four strains can be defined as new species.

**Table 1.**
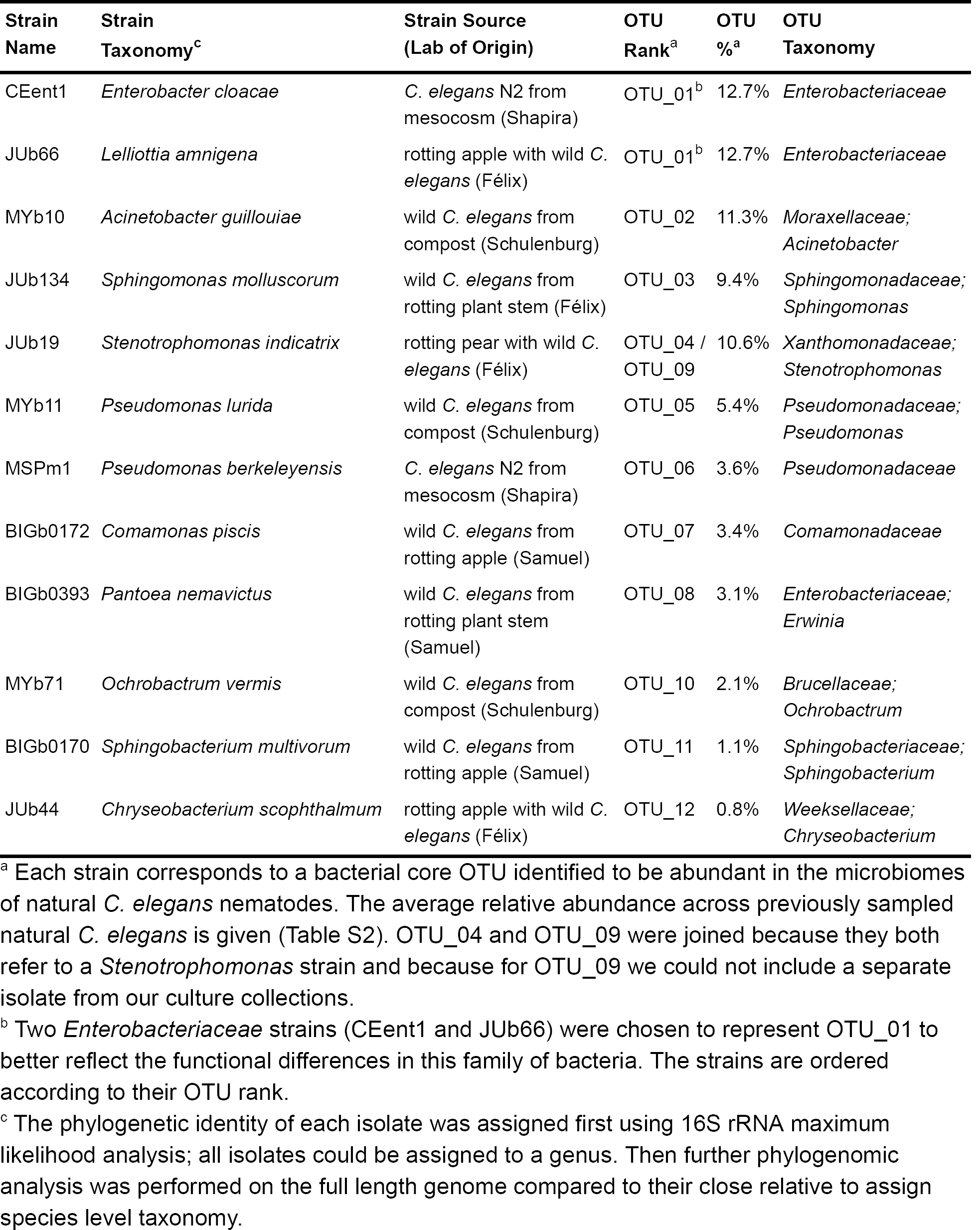
Members of the CeMbio v1.0 collection.

**Figure 1.**
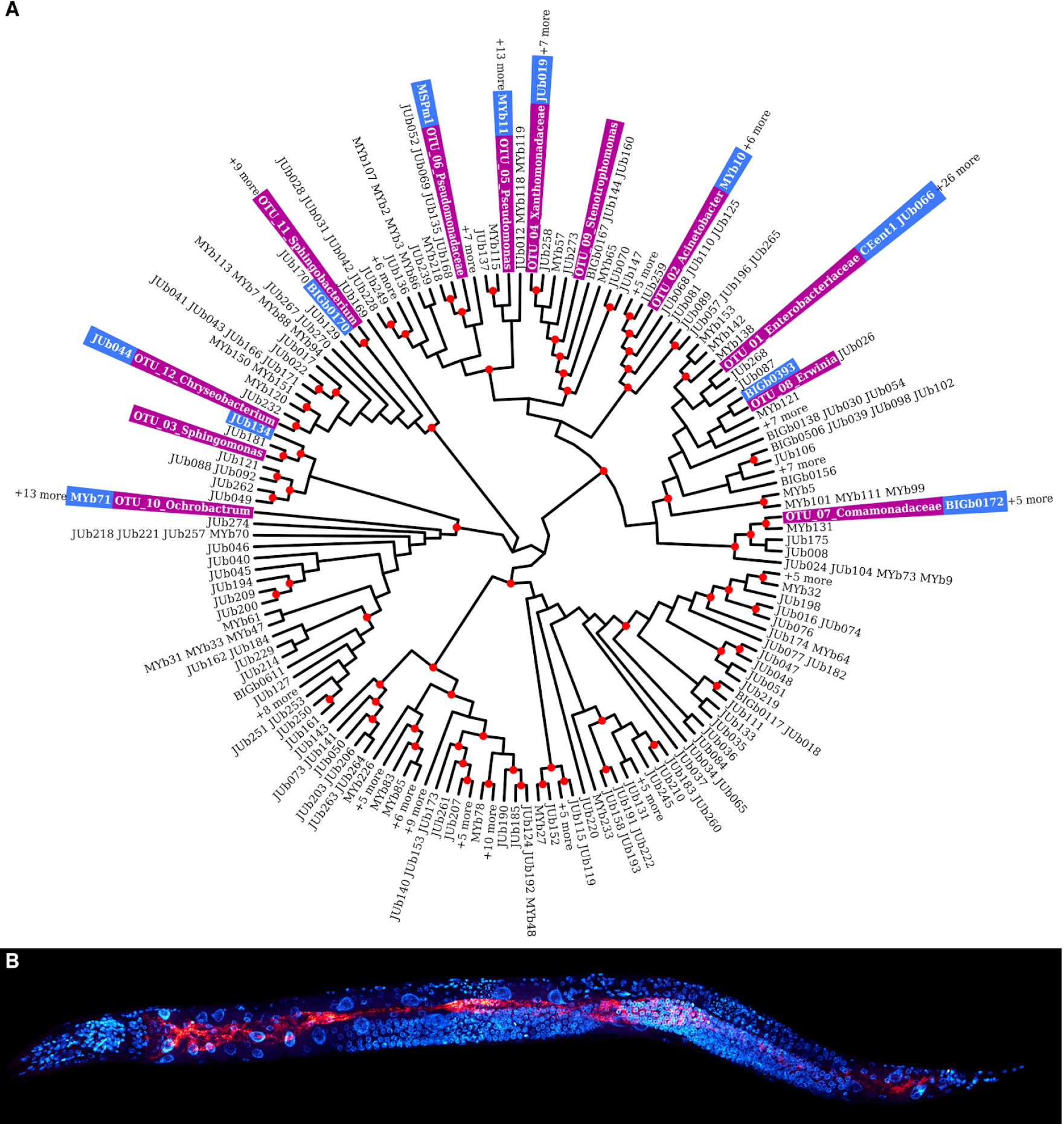
The CeMbio strains. (A) The CeMbio strains (blue) were selected based on a comparison of 510 cultured *C. elegans* microbiome bacteria with the 12 most common OTUs inferred by repeating our previous meta-analysis (purple, (Zhang *et al*. 2017)) with only the natural worm samples. The tree is based on a maximum-likelihood analysis using a TIM3e+R4 model and 10000 bootstraps. Nodes with bootstrap support > 75 % are denoted with a red dot. Some branches include several highly similar OTUs, as indicated (e.g., +6 more). (B) Fluorescence *in situ* hybridization of *C. elegans* N2 colonized with the CeMbio strains. Red: general bacterial probe EUB338; blue: DAPI.

For rapid PCR-based species identification, we designed diagnostic primers in genes that are unique to each isolate of the CeMbio resource. Unique genes were identified by reciprocal BLASTing (Altschul *et al*. 1990) of the respective genomes (see below). Primers targeting these unique genes were designed with primer3 (Untergasser *et al*. 2012) (Table S3), and subsequently assessed by PCR for strain specificity (Figure S1), using 0.2 µM of each primer and DreamTaq reagents (ThermoFisher Scientific) according to the manufacturer’s specifications with an annealing temperature of 60°C, and annealing and elongation times of 30 seconds each.

The bacteria were cryo-preserved in 15% glycerol/LB at −80°C to minimize laboratory adaptation. Growth and maintenance of the CeMbio strains can be achieved using identical methods as those for *E. coli* OP50. All CeMbio strains grow in Luria broth medium (LB; 10 g/l tryptone, 5 g/l yeast extract, 5 g/l NaCl, with or without 15 g/l agar) at 25 - 28°C (20 - 30°C possible) and reach stationary phase within 24 - 48 h (Figure S2). Other rich media such as TSB can be used, too. On LB-agar plates, most strains will produce single colonies after 24 - 48 h of incubation at 25°C. Slower-growing strains (e.g., JUb134) will yield visible single colonies only after 48 – 72 h, depending on inoculum size.

### CeMbio colonization experiments with *C. elegans*

We performed three independent experiments to assess the ability and dynamics of the CeMbio strains to colonize the nematode gut. The first experiment characterized colonization by each individual CeMbio strain separately, while the second and third experiments focused on colonization by the CeMbio community. The methods for these experiments are generally similar, yet deviated in particular aspects of the protocols, thereby allowing us to assess robustness of the results. In experiment 1, colonization was assessed by counting colony forming units (CFU) of bacteria isolated from nematodes. In experiments 2 and 3, colonization was characterized through CFU counts for the entire community and separately an analysis of the relative abundance of strains, inferred from 16S microbiome sequencing. The experiments were performed with the canonical *C. elegans* strains N2 (all experiments) and CB4856 (only experiment 2). Nematodes were maintained on nematode growth medium (NGM) seeded with a lawn of *Escherichia coli* OP50, as previously described (Stiernagle 2006). Below, we describe the methods used for each experiment.

#### Experiment 1

Experiment 1 assessed colonization levels by each CeMbio strains separately and as a community in *C. elegans* (N2) gut. The experiment generally followed the previously published protocol from the Samuel lab (Zhang *et al*. 2020) (protocols.io DOI: dx.doi.org/10.17504/protocols.io.rtzd6p6). Briefly, each CeMbio strain was grown individually in LB overnight at 25°C. Cultures were harvested by centrifugation, adjusted to a final optical density (OD, 600 nm) of 1 in PBS. Around 50 synchronized *C. elegans* stage 1 larvae (L1) were raised at 20°C on NGM in 6-well plates, each well inoculated with 60 µl bacteria. Nematodes and bacterial lawns were harvested after 72 h and 120 h with 600 μL of M9-T (M9 + 0.025% Triton X-100) and transferred to a sterile 96-well deep plate. To remove surface adherent bacteria, worms were washed five times with M9-T. After each washing step, worms were pelleted by centrifugation and aspiration of the supernatant using an aspiration manifold (V & P scientific, INC.). After the final wash, worms were left in 100 μl M9-T for 10 min in order to enhance digestion or defecation of any transient gut bacteria. 100 μl 10 mM levamisole solution was added to paralyze worms, followed by surface sterilization using 200 μl 4% bleach solution in M9 for 2 min. Thereafter, nematodes were washed twice with PBS to remove excess levamisole and bleach. 300 µl of the worms in PBS were combined with 1 mm sterilized garnet beads, followed by lysis in a Mixer Mill at 25 Hz for 5 min. Lysates were used directly for inference of the number of colony forming units (CFUs). CFU numbers were calculated by adapting a previously published protocol (Hazan *et al*. 2012). In short, a reference curve for microbiome abundance is generated that relates standardized CFU counts on plates to OD measurements of a corresponding culture in liquid. For both approaches, a dilution series was established for each CeMbio strain and then measured in parallel for the two methods in four replicates. The resulting reference curve was subsequently used to calculate CFU counts from OD measurements for the experimental samples.

#### Experiment 2

Experiment 2 served to assess colonization levels and composition of two *C. elegans* strains, N2 and CB4856, by the CeMbio community. It was based on the same protocols used for experiment 1 and included the following modifications. The CeMbio community inoculum was established by mixing equal volumes of the different bacterial strains, grown and processed as above. Worms were harvested after only 120 h. The obtained lysates were split in two and then either used directly for CFU inference (as above, File S2) or pelleted by centrifugation and frozen at −20°C for later microbiome analysis.

For microbiome analysis, DNA was extracted from frozen lysate pellets. The pellets were resuspended in 200 µl sterile PBS, 0.1 mm sterile zirconia/silica beads were added, and bacterial cells were further lysed in a Mixer Mill for 5 mins (25 Hz). 190 μl of the lysate was combined with 10 μl of 20 mg/ml proteinase K in a PCR plate and incubated in a MasterCycler ProS (Eppendorf) for 60 min at 60°C for digestion, followed by 15 min at 95°C to deactivate the proteinase. Barcoded amplicon sequencing was prepared according to the Earth Microbiome project (Caporaso 2018; Ul-Hasan *et al*. 2019) using the V4 region and sequenced by the Center for Metagenomics and Microbiome Research, Houston, Texas, USA.

#### Experiment 3

Experiment 3 used a slightly different approach to similarly study colonization of the *C. elegans* N2 strain by the CeMbio community in the Schulenburg lab. The CeMbio strains were grown individually in LB medium overnight at 28°C. Cultures were harvested by centrifugation, washed three times with PBS, and adjusted to a final OD_600_ of 5. The cultures were mixed in equal volumes to produce the CeMbio community inoculum. Synchronized *C. elegans* L1 animals were raised at 20°C on 6 cm plates containing either NGM or peptone-free NGM (PFM) seeded with 250 μl of the CeMbio inoculum. Nematodes and bacterial lawns were harvested after 48 h, 72 h, and 96 h with M9-T. Surface-adherent bacteria were removed using a modification of a previously described method (Dirksen *et al*. 2016; Papkou *et al*. 2019) (Figure S3). Briefly, suspended worms were placed onto the top of pipette tips containing a 10 µm filter and repeatedly incubated with washing solutions: 2x 3 min of M9-T with 25 mM tetramizole hydrochloride (to anesthetize worms and prevent subsequent bleach intake); 1x 4 min of M9-T with 2% bleach (equal volumes of 12% sodium hypochlorite and 5 N NaOH, (Stiernagle 2006)); 2x 3 min of M9-T to remove the bleach. After each washing step, the solution was removed by centrifugation of the tip box. The washed worms were pelleted by centrifugation and either frozen at −20°C for microbiome analysis or subjected to immediate CFU extraction. For the latter, ten nematodes were transferred to a 2 ml tube containing 100 µl M9-T and 10–20 1 mm zirconium beads, followed by sample homogenization using a Geno/Grinder 2000 (SPEX SamplePrep, Metuchen, USA) at 1500 strokes/min for 3 min. The homogenate was serially diluted in M9-T and each dilution was plated on LB-agar in triplicates. After 48 h of incubation at 25°C, the plates were imaged and appropriate dilutions counted.

For the microbiome analysis, DNA was isolated from frozen surface-sterilized worm samples or frozen lawn pellets, resuspended in buffer T1 from the NucleoSpin® Tissue Kit (Macherey & Nagel), and processed with the additional steps described in the “Support protocol for bacteria” following the manufacturer’s instructions. Barcoded amplicon sequencing of the V3-V4 region of the bacterial 16S rRNA gene was carried out by the Institute for Clinical Molecular Biology, Kiel, Germany, using Illumina MiSeq technology.

### Fluorescence *in situ* hybridization

Fluorescence *in situ* hybridization of the CeMbio strains colonizing the *C. elegans* gut (Figure 1B) was performed as previously described (Dirksen *et al*. 2016; Yang *et al*. 2019).

### Microbiome data analysis

Sequencing data was prepared for subsequent statistical analysis by first removing adapter and primer sequences with cutadapt (Martin 2011). Amplicon sequence variants (ASVs) were inferred using the R package *dada2* (Callahan *et al*. 2016) with default parameters except for the following settings: sequence truncation length forward/reverse: 250/200 (longest expected amplicon for V3-V5: 428 nt, for V4-V5: 250nt); taxonomic assignment with silva trainset release 132 (Quast *et al*. 2013); species assignment with a custom reference set of genome-derived 16S sequence variants of the CeMbio strains. The ASV read counts were normalized by the 16S gene copy numbers of the corresponding bacterial strains as predicted by the genome assemblies prior to analysis. The statistical analysis of the ASV data was performed in R using the following packages: *DECIPHER* (Wright 2016), *phyloseq* (McMurdie and Holmes 2013), *DESeq2* (Love *et al*. 2014), *vegan* (Oksanen *et al*. 2019), *ggplot2*.

### Developmental Timing

To assess the influence of CeMbio strains on *C. elegans* development, nematodes were raised on single and mixture lawns and the number of adults over time was counted, following a previously published protocol (Samuel *et al*. 2016). In brief, the 12 CeMbio bacteria were grown in LB at 25°C overnight; *E. coli* OP50 was also assayed for comparison, yet grown at 37°C. Bacteria were concentrated and seeded (50 μl) into 6-well NGM plates in duplicate. Plates were dried and incubated overnight at 25°C before adding around 100 synchronized L1 worms to the wells containing either a single bacterial strain or the CeMbio mixture. Adult animals were scored on an hourly basis from 46-60 h post L1 stage.

### CeMbio genome sequences and metabolic network reconstructions

Bacterial genomes were sequenced using short (Illumina Nextera XT for CEent1, MYb10, MYb11, MYb71, and MSPm1; and all remaining isolates with Illumina Miseq v3) and long read (PacBio SMRT; all isolates) sequencing. Short read Illumina reads were preprocessed with fastq_illumina_filter 0.1 (--keep N -vv) and prinseq-lite 0.20.4 (-min_len 20 -ns_max_n 8 -min_qual_mean 15 -trim_qual_left 12 -trim_qual_right 12) (Schmieder and Edwards 2011), followed by barcode demultiplexing and filtering of the long reads with lima 1.8 (--peek-guess --split-bam-named) (https://github.com/PacificBiosciences/barcoding). The genomes were assembled by combining short and long reads in a hybrid approach, using the following programs: SPAdes v3.13.1 (Bankevich *et al*. 2012), Canu 1.8 (Koren *et al*. 2017), MaSuRCA 3.3.4 (Zimin *et al*. 2013), and the Unicycler pipeline 0.4.8 (Wick *et al*. 2017). Long reads correction was achieved with LoRDEC 0.6 (Salmela and Rivals 2014), proovread 2.14.1 (Hackl *et al*. 2014), and Canu 1.8 (detailed script with all program calls and parameters is available in the supplement). The quality of genome assemblies was assessed with QUAST 5.0 (Mikheenko *et al*. 2018). The final genomes were derived after assessing the quality by coverage vs. length plots and by removing low quality contigs with <500bp and <5 coverage (Douglass *et al*. 2019).

The genome assemblies served as input for the reconstruction of metabolic networks, using gapseq 1.0 (Zimmermann *et al*. 2020b). In detail, pathways and transporters were predicted by *gapseq find* (-b 150), and the draft network was created by *gapseq draft* (-u 150 -l 50 -a 1). Network gaps were filled with *gapseq fill* (-b 50). Metabolic networks were thus represented by genome-scale metabolic models and combined with flux balance analysis (Orth *et al*. 2010), in order to predict growth rates under specified conditions. Gap filling was focused on ensuring bacterial growth in LB medium, which is known to support the growth of all CeMbio organisms in experiments. The metabolic networks were further improved with gapseq 1.0 by integrating experimental data derived from EcoPlate assays (Biolog, Inc, USA), in which the reduction of a colorimetric tetrazolium dye indicates microbial metabolic activity on selected carbon sources (Bochner 2009), thereby providing empirical information on the metabolic competences of the CeMbio strains. The metabolic network models were subsequently used to predict carbon source utilization by the CeMbio strains, based on flux balance analysis with the recycling of electron carriers (quinones, NADH) as objective function. An organism was predicted to be able to use a certain compound if electron carriers could be recycled under conditions of a minimal medium including this compound as sole energy and carbon source. The inferred metabolic network models are available in the supplement (SBML format; Files S2 and S3).

### CeMbio phylogenomic reconstructions

Genome-scale phylogenies were calculated using GToTree V1.4.11 (Lee 2019). Each step of the pipeline was used with the default parameters. In brief, for each CeMbio strain, we downloaded NCBI RefSeq assemblies belonging either to the same genus or the same family depending on the number of published related genomes that were available. For genera with a large amount of genomes available, such as *Enterobacteriaceae* and *Pseudomonas*, we downloaded only genomes annotated as complete for representatives. For the less represented genera, we included partial assemblies. Genomes without annotation were scanned for CDS using prodigal (Hyatt *et al*. 2010), then genes were scanned for single-copy marker genes using HMMER3 (Eddy 2011), genomes with less than 10% of single marker gene redundancy were kept. Then single-copy marker genes were aligned using MUSCLE (Edgar 2004), trimmed with TrimAl (Capella-Gutiérrez *et al*. 2009) in order to keep sequence overlap and finally phylogenetic tree were calculated using FastTree 2 (Price *et al*. 2010). An Alphaproteobacteria, *Bradyrhizobium diazoefficiens* (GCF_000011365.1), was arbitrarily chosen as an outgroup for all trees. Taxonomy was edited on the tree using Taxonkit (Shen and Xiong 2019). For additional details on phylogenomic reconstructions, the phylogenomic tree as well as the code used for the analysis see File S3.

To evaluate the taxonomic affiliation of each CeMbio bacteria, we compared 16S rRNA phylogeny and phylogenomic reconstructions, then estimated the relatedness of each genome to their close phylogenomic relative using average nucleotide identity (ANI). ANI was calculated using a script available from the Enve-omics package (Rodriguez-R and Konstantinidis 2016). Bacteria with closely related genomes were compared and we used a ANI of 94-96% for the species cutoff, as described in previous studies (Konstantinidis and Tiedje 2005; Richter and Rosselló-Móra 2009). In the case where the 16S rRNA phylogenetic reconstruction provided a closely related named species that were not sequenced and no genomes were closely related to the CeMbio bacteria we relied on the 16S ribosomal phylogeny for strain naming purposes.

### Resource Sharing and Data Availability

To facilitate broad distribution of CeMbio strains, all isolates are available from the *Caenorhabditis* Genetics Center (CGC, cgc.umn.edu; search for ‘CeMbio’ in strain descriptions). Whole genome sequencing data (PRJNA624308) and microbiome 16S sequencing data are publically available [PRJEB37101 (experiment 2), PRJEB37035 (experiment 3)]. All experimental data is provided in the supplement (see File S2) and have been uploaded to the GSA Figshare Portal. Additional information on laboratory methods can be found at the CeMbio Wiki (www.CeMbio.uni.kiel.de).

## RESULTS

### Overview of the CeMbio resource

We here established an ecologically informed model *C. elegans* microbiome (CeMbio) based on the following key criteria: (i) the chosen community should resemble the broad taxonomic diversity of the natural *C. elegans* microbiome as closely as possible; (ii) it should be ecologically meaningful and thus originate from natural *C. elegans* or at least its natural habitat; and (iii) it should include bacteria that are easy to grow and maintain on a standard medium, thus facilitating experiments in different fields of biological research.

As a first step, we repeated our previous microbiome analysis using only the natural *C. elegans* samples (Zhang *et al*. 2017), in order to identify the most abundant worm-associated bacterial taxa (Table S2). A total of 12 OTUs was consistently present in natural *C. elegans*, regardless of origin, isolating laboratory, or other covariates. This set of bacterial OTUs is likely to represent an ecologically relevant part of the *C. elegans* microbiome and also covers a substantial proportion of its natural diversity (∼63% of the microbiome diversity found in natural *C. elegans* isolates).

Thereafter, we identified the best 16S rRNA sequence matches between each of these OTUs and the bacterial isolates from our culture collections of microorganisms from natural *C. elegans* or *C. elegans*-containing substrates (available in the Félix, Samuel, Schulenburg, and Shapira labs; Table 1). The final selection was prioritized according to the source sample for each isolate (*C. elegans* animal was preferred over habitat), sequence identity (exact match preferred) and body of existing knowledge (priority to those that had already been characterized in more detail). For the OTUs without exact matches among the isolates, we chose the strain with the lowest phylogenetic distance from the OTU. Using this approach, we identified sequence-identical isolates or closely related strains for all but one of the 12 most abundant OTUs (Figure 1, File S1). The missing OTU referred to a *Stenotrophomonas* strain, for which we do not have a closely related isolate in our collections. However, another Stenotrophomonas isolate was included as a perfect match for a different OTU (i.e., OTU_04, Figure 1). We further included two isolates of the most abundant OTU from the Enterobacteriaceae (i.e., OTU_01).

Based on the above analyses, we selected 12 isolates to constitute the CeMbio resource. For these 12 isolates, we developed diagnostic PCR primers, thus allowing their identification within the community (Figure S1). These 12 isolates can be maintained on standard LB and NGM medium (Figure S2).

### Individual CeMbio strains effectively colonize the *C. elegans* intestine

In experiment 1, we determined whether and to what extent each of the CeMbio bacteria are able to colonize the *C. elegans* intestine. Based on previous studies with other non-pathogenic bacteria, we expected colonization of the *C. elegans* intestine during early adulthood (Dirksen *et al*. 2016; Vega and Gore 2017). To examine this directly with the CeMbio strains, synchronized L1 animals were exposed to each bacterium for 72 h and 120 h, those time points respectively corresponding to the first and third day of adulthood for N2 nematodes growing on OP50 *E. coli*. All strains were able to colonize the intestines of *C. elegans*, in overall agreement with previous studies, where some of these strains had been examined (Montalvo-Katz *et al*. 2013; Berg *et al*. 2016a, 2019; Dirksen *et al*. 2016; Zimmermann *et al*. 2020a). However, the extent and persistence of colonization over time varied among strains. Eleven of the twelve strains exhibited colonization levels that increased 10-fold over time (from an average of ∼500 CFUs/worm at 72 h to ∼10000 CFUs/worm at 120 h; Figure 2). Only the *Sphingomonas* JUb134 exhibited a slow colonization increase overtime (3-fold), which could be linked to its long doubling time. At the other end of the spectrum, *Ochrobactrum sp*. MYb71 showed a dramatic 38-fold increase in colonization between the timepoints. In general, the strains could be organized into three groups: (1) the low colonizers, including all three *Enterobacteriaceae* (*Pantoea* BIGb0393, *Enterobacter* CEent1, *Lelliottia* JUb66) and *Acinetobacter* sp. MYb10; (2) the intermediate colonizers, *Sphingomonas* JUb134, *Comamonas* sp. BIGb0172, *Sphingobacterium* sp. BIGb0170 and *Pseudomonas* sp. MYb11; and (3) the high colonizers, *Ochrobactrum sp*. MYb71, *Chryseobacterium* sp. JUb44, *Pseudomonas* sp. MSPm1, and *Stenotrophomonas* sp. JUb19. We conclude that the selected CeMbio strains are all able to establish themselves in the gut of *C. elegans*, providing ample opportunity for direct interactions between microbe and worm.

**Figure 2.**
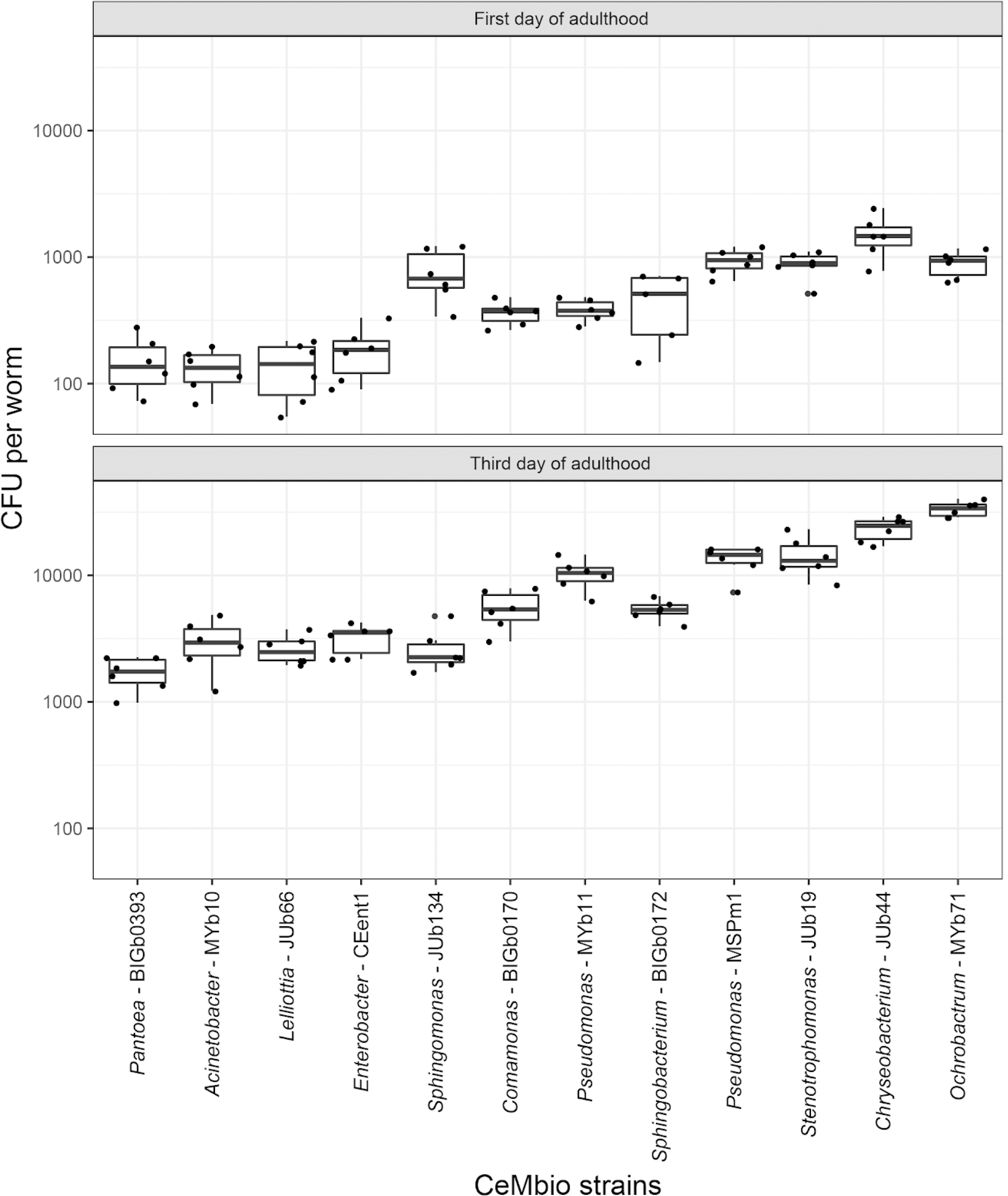
Colonization levels of *C. elegans* gut by each CeMbio strain alone. Colony forming units (CFUs) of each CeMbio strain in *C. elegans* gut (N2) were measured at 72 h or 120 h post L1 larvae. At least six biological replicates were performed for each condition. These results are from colonization experiment 1.

### All CeMbio strains colonize the worm gut when they are part of a community

In experiments 2 and 3, we next assessed whether the individual CeMbio strains colonize *C. elegans* while being part of a community. We performed two independent experiments that were performed in different labs and varied in culture media, sampling time points, and exact processing protocols (see methods for more details). Overall, the experiments demonstrate that (i) a single *C. elegans* adult is consistently colonized by at least 1,000 and usually more than 10,000 bacteria (Figure 3C, 4F), (ii) all CeMbio strains can establish themselves in the worm gut as community members (Figure 3B, 4B, 4C), (iii) the exact colonization dynamics depend on bacterial strain, time, and culture medium (Figure 3B, 3E, 4B, 4C, 4E), and (iv) the *C. elegans*-associated community is clearly distinct in composition and diversity from the corresponding lawn community on the Agar plates (Figure 3B, 3D, 3E, 4B-4E).

**Figure 3.**
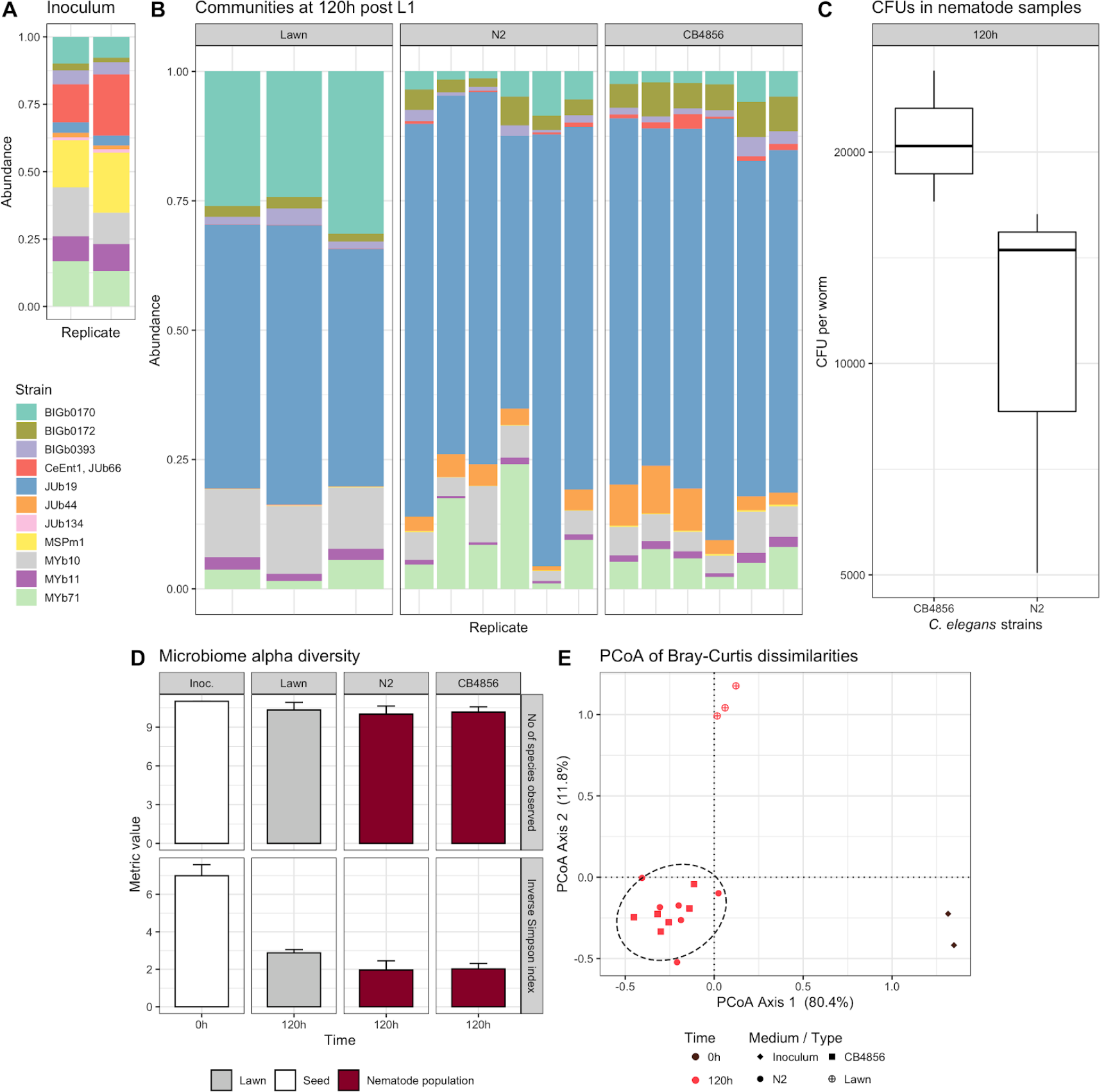
Colonization of N2 and CB4856 *C. elegans* strains by the CeMbio community. (A) Proportion of reads in the initial community assembly used as inoculum for the lawns. (B) Proportion of reads in the *C. elegans* strains N2 and CB4856 and the corresponding lawn samples. (C) Colony forming units (CFUs) of the CeMbio community isolated from N2 and CB4856 nematodes. (D) Mean observed number of CeMbio members (top) and Inverse Simpson Index (bottom) with standard deviation, indicating richness and diversity of the bacterial communities in N2 and CB4856 worms. (E) Principle coordinate analysis of Bray-Curtis dissimilarities of the microbial communities of nematode and lawn samples with an ellipse representing the 95% confidence interval of the nematode samples. These results are from colonization experiment 2.

The CeMbio strains were generally able to persist as a community both on plates and in nematodes. All 12 strains could be detected in the lawns of both NGM and PFM plates across the two independent experiments, even though some strains were present at very low levels (Figure 3B, 4B, 4C). Using improved protocols for nematode surface sterilization (Supplementary Figure 3; see Methods), we reliably detected all CeMbio strains as colonizers of *C. elegans* guts when part of a community despite some again appearing only at very low levels (Figure 3B, 4B, 4C). Three CeMbio strains were generally highly abundant in worms, including the *Sphingobacterium* sp. BIGb170, the *Stenotrophomonas* sp. JUb19, and the *Ochrobactrum* MYb71. While BIGb170 and JUb19 were more abundant in NGM-raised worms, MYb71 showed higher prevalence in the PFM-grown worms, with increasing proportions at the later time points. The abundance of other CeMbio members showed more variation dependent on experiment, medium, and time point. For example, the *Pantoea* BIGb393 only reached high abundance in worms from PFM plates at the latest time point, while the two *Pseudomonas* strains (MSPm1 and MYb11) were more abundant in worms from PFM rather than NGM plates, especially at the two early time points. Our results further highlight that colonization behavior within the community differs from that of individual bacteria. This is most apparent for the *Chryseobacterium* JUb44, which reaches high frequencies in worms when alone but low abundances when part of the community (Figures 2, 3B, 4B, 4C). Similarly, the *Pseudomonas* strains (MSPm1 and MYb11) are comparatively good colonizers when alone, while reaching only lower abundances within the community, particularly on NGM.

The *C. elegans* microbiome is generally very distinct from the corresponding microbial environment on the plates. This difference is most obvious in the PCoA analyses (Figure 3E, 4E), but also in the relative abundances (Figures 3B, 4B, 4C) and in experiment 3 in the observed diversities (Figure 4D). It is worth noting that the bacterial mixture used to inoculate the plates is best maintained on PFM (Figure 4A-4C, 4E), which does not support bacterial growth and thus appears to enhance experimental control of the source community.

**Figure 4.**
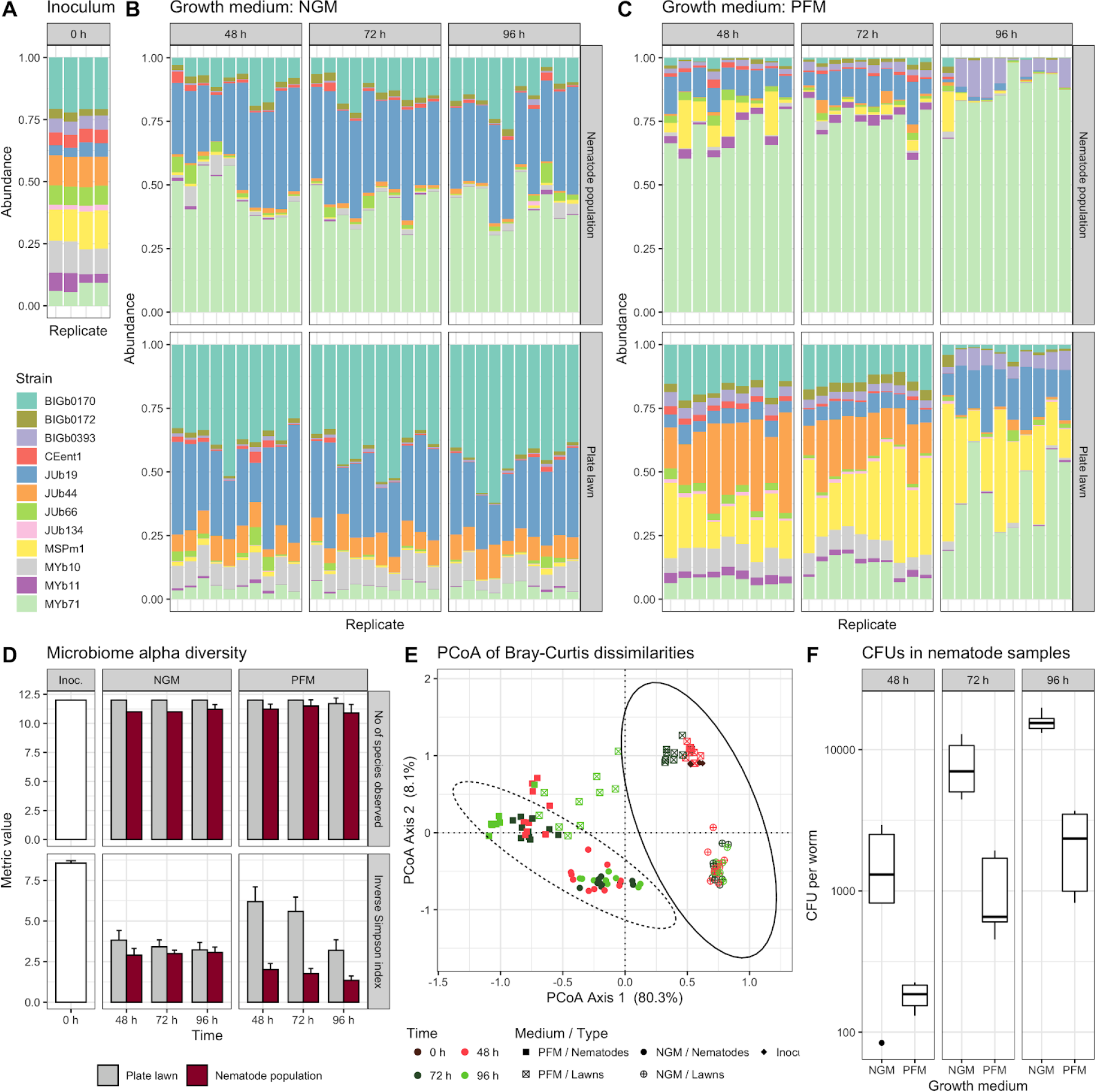
Colonization of *C. elegans* gut by the CeMbio community under different plating conditions. (A) Proportion of reads in the initial community assembly used as inoculum for the lawns. (B) Proportion of reads in NGM worm and lawn samples. (C) Proportion of reads in PFM worm and lawn samples. (D) Mean observed number of CeMbio members (top) and Inverse Simpson Index (bottom) with standard deviation indicating richness and diversity of the communities. (E) Principle coordinate analysis of Bray-Curtis dissimilarities of the microbial communities in nematode and lawn samples with ellipses representing the 95% confidence intervals of the nematode (dashed) and lawn (solid) samples. (F) Colony forming units (CFUs) of the CeMbio community in single nematodes. These results are from colonization experiment 3.

Using DESeq2 for differential abundance analysis (File S2), we found three CeMbio strains to be consistently enriched in nematodes relative to lawns: *Ochrobactrum* sp. MYb71 was always enriched regardless of medium or time point within a range of 8.8 - 20 fold. *Enterobacter* CEent1 was enriched regardless of medium in the adult stages (72 h and 96 h) within a range of 1.8 - 6.1 fold. *Lelliottia* sp. JUb66 was consistently enriched in PFM-raised worms (1.5 - 4.8 fold); although a similar consistent enrichment in NGM-raised worms by 1.7 - 3.2 fold is likely, it was statistically significant only for the 72 h time point. Other strains were enriched in a context-dependent manner. For example, *Pantoea* sp. BIGb0393 was either not enriched or even decreased (minimum: 0.37 fold) under all conditions, except in late PFM-raised worms, where it was strongly enriched (maximum: 8.0 fold) compared to its abundance in the lawns. JUb19, the second most abundant strain in worms, is either not or only minimally enriched (1.1 - 1.3 fold) on NGM, while being moderately enriched in PFM-raised worms for L4 larvae and early adults (48 h and 72 h; 3.7 and 2.4 fold, respectively), before being strongly depleted at 96 h (0.25 fold). Other strains are generally depleted in worms compared to lawns, most strongly among them JUb134 (0.076 - 0.0082 fold).

Taken together, these results suggest that the CeMbio community contains a combination of strong general colonizers (*Ochrobactrum* sp. MYb71 and two of the Enterobacteriaceae strains, CEent1, JUb66) and context-dependent colonizers (*Pantoea* sp. BIGb0393 and *Stenotrophomonas* sp. JUb19). *C. elegans* can be colonized by the CeMbio community under different conditions, thus enabling new research on the effect of the microbiome on nematode biology.

### CeMbio strains and community vary in impact on host growth rates

To illustrate the potential for the CeMbio strains to influence *C. elegans* biology, we characterized its phenotypic impact on nematode development rates. Two *C. elegans* strains (N2 and CB4856) were raised on single bacteria and the community from L1s and populations were followed over time for maturation to adulthood (Figure 5). In comparison to growth on *E. coli* OP50, worms developed faster on all single bacteria with three exceptions: on *Ochrobactrum* sp. MYb71, they developed at a similar pace, while on *Chryseobacterium* sp. JUb44 and *Sphingobacterium* sp. BIGb0170, nematodes developed at a significantly slower pace. This is consistent with previous studies that indicate slower growth of worms on Bacteroidetes strains like JUb44 and BIGb0170 (Samuel *et al*. 2016). Notably, the CeMbio community as a whole significantly enhanced growth rates for both host strains compared to growth on *E. coli* OP50. This suggests that the community may contain emergent properties that are produced as a result of interactions between the microbes that promote host development.

**Figure 5.**
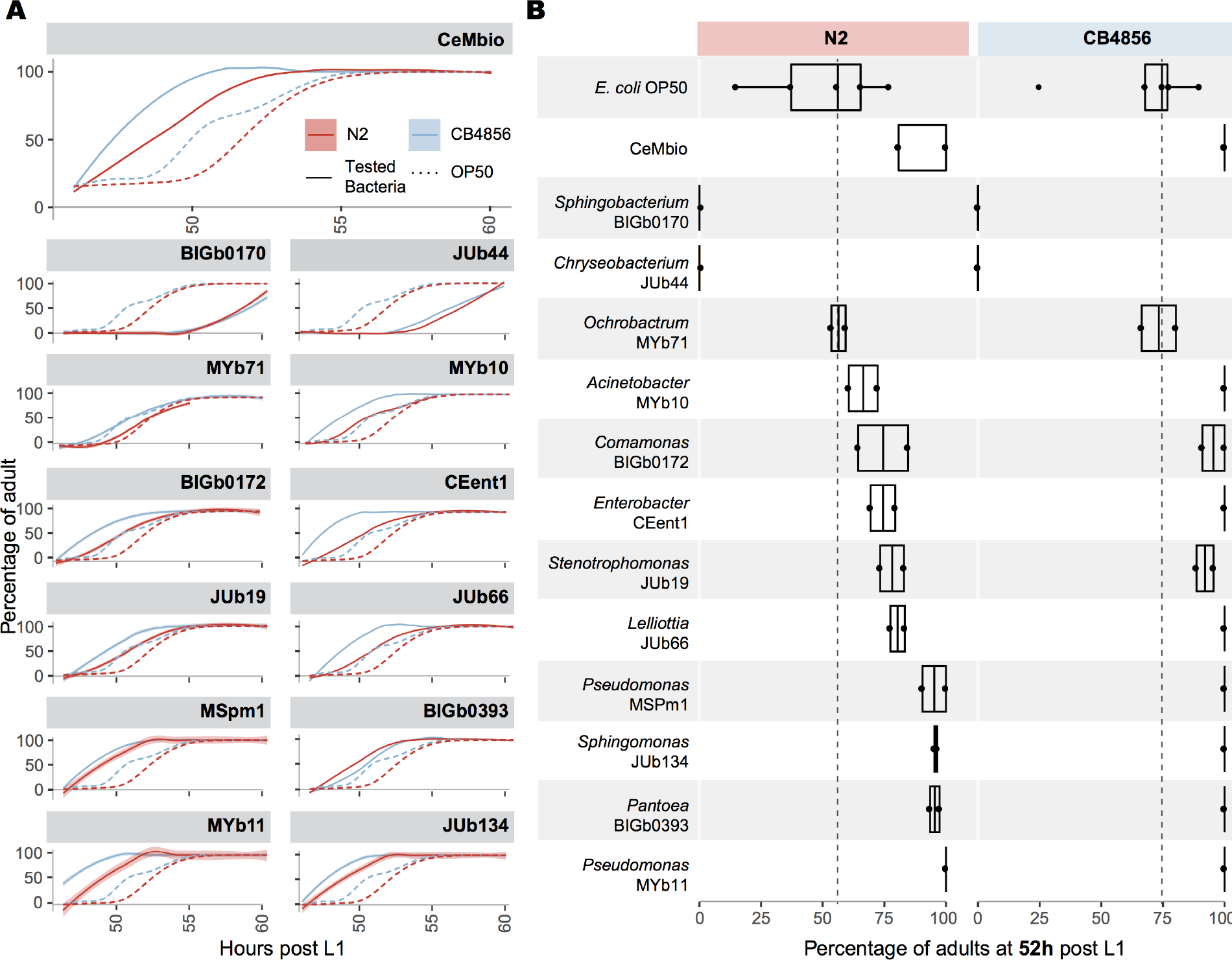
Effect of the CeMbio community and individual bacteria on *C. elegans* growth rates. (A) Developmental speed, represented by the number of adults counted on an hourly basis, when N2 and CB4856 nematodes are raised on the CeMbio mixture or the individual bacteria. Continuous lines indicate CeMbio bacteria or mixture; the dotted line the *E*.*coli* OP50 control. (B) Developmental timing snapshot at 52 h post L1. The black dotted line represents the median number of adult worms on *E. coli* at that time point where roughly 50% of the N2 population reached adulthood (n=50-100 animal/replicate).

Our results also suggest a range of host-bacteria interaction types for the CeMbio members that may have both positive and negative impacts on the host. For example, JUb44 are efficient colonizers individually, yet slow down the developmental rate of its host, possibly indicating a negative effect of the bacteria on *C. elegans*. Though as part of the community this negative impact is mitigated, and also consistent with previous studies where growth rates improved with when JUb44 lawns included as little as 5% of a Proteobacteria strain (Samuel *et al*. 2016). Good colonizers, like *Stenotrophomonas* sp. JUb19, also have a positive effect on host developmental speed, possibly indicating a beneficial association. Other bacteria such as *Pantoea* sp. BIGb0393 increase developmental speed yet are bad colonizers; they may thus represent a good source of nutrition for the worm despite lacking a more intimate interaction with its host. When colonized by the bacterial community, the apparently negative effect of certain single bacteria seems to be counterbalanced by the presence of other bacteria.

### Using whole genome sequences to reconstruct strain phylogenies

To add to the value of this resource for the community and spur future in-depth analysis of *C. elegans*-microbiome interactions, we sequenced the genome of each of the 12 isolates. Using a combination of short and long reads and a hybrid approach for genome assembly, complete genome sequences were obtained for all CeMbio members (Table 2).

**Table 2.** Characteristics of the CeMbio genomes^a^.

| Strain | Genus | Size | GC | Contigs | rRNA | tRNA | CMP | CDS | UG | CRISPR | VG | RG |
| --- | --- | --- | --- | --- | --- | --- | --- | --- | --- | --- | --- | --- |
| CEent1 | *Enterobacter* | 4.8 | 55.3 | 1 | 8 | 89 | 98 | 4458 | 2787 | 0 | 38 | 4 |
| JUb66 | *Lelliotta* | 4.6 | 52.9 | 1 | 7 | 84 | 98 | 4207 | 2684 | 3 | 33 | 3 |
| BIGb0393 | *Pantoea* | 5.2 | 54.6 | 2 | 7 | 82 | 98 | 4667 | 2540 | 0 | 21 | 1 |
| MYb10 | *Acinetobacter* | 4.6 | 38.3 | 1 | 7 | 82 | 98 | 4244 | 1611 | 1 | 2 | 2 |
| JUb19 | *Stenotrophomonas* | 4.6 | 66.3 | 2 | 4 | 78 | 96.6 | 4079 | 1625 | 0 | 13 | 5 |
| MYb11 | *Pseudomonas* | 6.1 | 60.8 | 1 | 5 | 66 | 100 | 5456 | 2248 | 0 | 78 | 1 |
| MSPm1 | *Pseudomonas* | 5.7 | 62.4 | 2 | 4 | 70 | 100 | 5159 | 2017 | 1 | 83 | 0 |
| BIGb0172 | *Comamonas* | 5.2 | 62.6 | 1 | 6 | 81 | 95.9 | 4595 | 1719 | 0 | 3 | 1 |
| MYb71 | *Ochrobactrum* | 5.4 | 55.9 | 3 | 4 | 60 | 97.3 | 5191 | 1814 | 0 | 16 | 2 |
| JUb134 | *Sphingomonas* | 4.1 | 67.5 | 4 | 3 | 65 | 96.6 | 3816 | 1292 | 0 | 2 | 0 |
| BIGb0170 | *Sphingobacterium* | 6.4 | 39.9 | 1 | 7 | 85 | 93.2 | 5391 | 1362 | 0 | 1 | 0 |
| JUb44 | *Chryseobacterium* | 4.7 | 33.6 | 1 | 7 | 80 | 94.6 | 4199 | 1173 | 1 | 1 | 2 |

<sup>a</sup> The table highlights predicted key features of the genome assemblies. Size: genome size in Mb; GC: percent GC content; Contigs: total no. of contigs of the assembly; rRNA: no. of genomic rRNA as predicted by RNAmmer (Lagesen et al., 2007); tRNA: Number of genomic tRNA as predicted by ARAGORN (Laslett and Canback, 2004); CMP: percent genome completeness as predicted by BUSCO (Simão et al., 2015); CDS: Number of coding sequences as predicted by PROKKA (Seemann, 2014); UG: Number of unique genes per assembly (PROKKA); CRISPR: Number of genomic CRISPR genes (PROKKA); VG, RG: Number of virulence and resistance genes as predicted by Abricate (Seemann T, Github https://github.com/tseemann/abicate) using the Virulence Factor Database (Chen et al., 2016) or the NCBI Bacterial Antimicrobial Resistance Reference Gene Database (Feldgarden et al., 2019), respectively.

Complete genomes allowed us to perform Phylogenomic analysis and assign an accurate species level phylogeny for 10 out 12 CeMbio species. In brief, in the case of BIGb0172 and JUb134, not enough published genomes were available to reconstruct a reliable phylogenomic tree. We then used the 16S rRNA phylogenies (File S1) to classify those organisms as *Comamonas piscis* for BIGb0172 and *Sphingomonas molluscorum* for JUb134. For the bacteria MYb11, Myb10 and JUb66, phylogenomic reconstruction yielded similar trees as their respective 16S rRNA phylogenetic counterparts and we confirmed their assignment as *Pseudomonas lurida, Acinetobacter guillouiae* and *Lelliottia amnigena*. The bacteria CeEnt1, BIGb0170 and JUb19 could be attributed more accurately to a single species, respectively *Enterobacter cloacae, Sphingobacterium multivorum* and *Stenotrophomonas indicatrix*. Our phylogenomic analysis further indicated that BIGb0393, MYb71, JUb44 and MSPm1 are new species of, respectively, the *Pantoea, Ochrobactrum, Chryseobacterium* and *Pseudomonas* genera (see File S3; Table 1).

### Whole genome sequences reveal diverse metabolic competences of the CeMbio strains

Based on these genomes, we determined the presence or absence of different metabolic pathways and the overall metabolic network for each CeMbio strain. We found that the metabolic potential of the CeMbio bacteria ranges from 186 pathways present in the *Chryseobacterium* sp. JUb44 to 389 pathways in *Enterobacter* sp. CEent1 (Figure 6A). Overall, common pathways are present in similar abundance across the genomes while more unique pathways are more unevenly distributed. Both *Pseudomonas* (MSpm1 and MYb11) and *Enterobacteriaceae* (*Pantoea* sp. BIGb0393, *Enterobacter* sp. CEent1, *Lelliottia* sp. JUb66) strains have overall more pathways and more unique pathways, while *Chryseobacterium* sp. JUb44 and *Comamonas* sp. BIGb0170 have fewer predicted pathways overall and also fewer unique pathways (Figure 6A). A principal component analysis of the metabolic potential of the 12 bacteria shows a clustering related to taxonomy, with distinct groupings for the *Enterobacteriaceae, Bacteroidetes* and *Pseudomonas* (Figure 6B).

**Figure 6.**
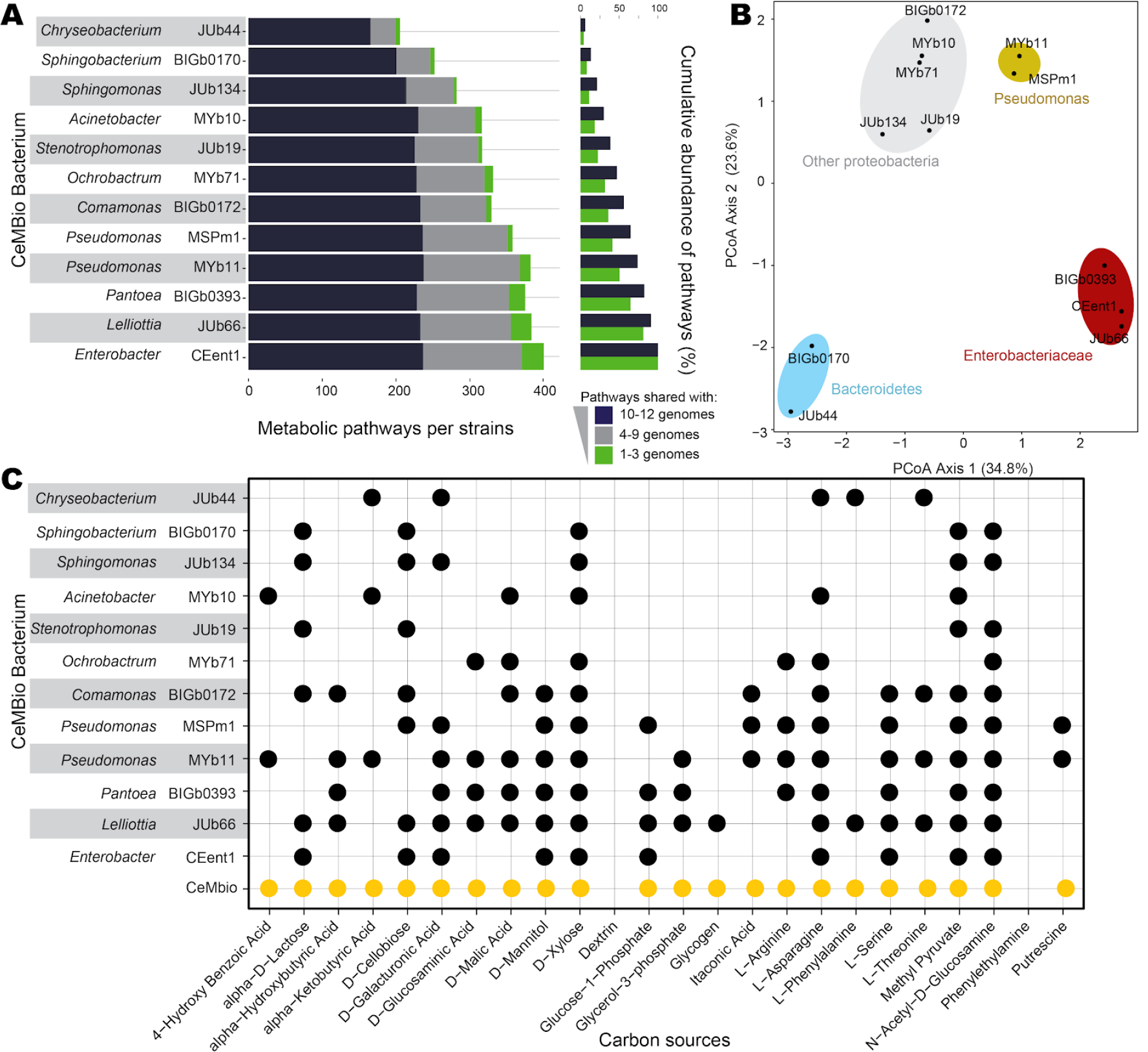
Comparison of metabolic pathways between the 12 CeMbio strains. **(A)** Distribution of unique and shared metabolic pathways across the 12 CeMbio members. Pathways are categorized from the most commonly found (present in 10 to 12 genomes) to unique pathways (present in 1 to 3 genomes). **(B)** Principal component analysis of the metabolic profiles of the 12 CeMbio members. **(C)** Summary of carbon source utilization for each CeMbio strain as inferred from the genome-scale metabolic models, additionally trained with Biolog EcoPlate plate data. The carbon source utilization by the whole CeMbio community is highlighted in yellow.

As an illustration of the potential for interactions between the strains, we subsequently used metabolic modeling to predict the range of carbon sources that each strain can utilize and compared it with experimental results obtained from Biolog microarrays (File S2). We found that 66% of the experimental data on carbon utilization were consistent with the predictions, which was similar to our previous study (Zimmermann *et al*. 2020a). In addition, the experimental data were subsequently used to further optimize the metabolic models (File S4). These adjustments led to an overlap of 99% between the model predictions of usable carbon sources and the Biolog results. In general, the metabolic network model analysis indicated variation in the metabolic competences of the CeMbio strains. Almost all strains can utilize and likely would compete for specific carbon sources like pyruvate, xylose, and N-acetyl-D-glucosamine. However, the bacteria varied in their abilities to process other carbon sources such as glycogen, phenylalanine, or benzoate (Figure 6C). Together, this type of metabolic niche-partitioning may explain some of the consistently identified community structure that we have observed when colonizing the *C. elegans* gut versus what is observed in the lawn.

Metabolic network modelling further indicates that the CeMbio community can provide metabolites important for *C. elegans* growth (File S4). This assessment is in general agreement with previous work. For example, a strain of the soil bacterium *Comamonas* was shown to provide vitamin B12 to the worm, which in turn influences development and fertility through the methionine/SAdenosylmethionine cycle while it also processes propionic acid, thereby removing its toxic effects (Watson *et al*. 2014, 2016). Interestingly, the two CeMbio members MYb11 and MYb71, usually enriched in the *C. elegans* microbiome (Dirksen *et al*. 2016; Zhang *et al*. 2017; Zimmermann *et al*. 2020a; Johnke *et al*. 2020); see also below), also possess the pathways for vitamin B12 production (Zimmermann *et al*. 2020a) and could thus influence similar *C. elegans* characteristics as the *Comamonas* strain. Together, these studies illustrate how CeMbio strain combinations and underlying genetic potential can facilitate interrogation of ecologically relevant influence of microbial metabolites on a wide range of life history characteristics and aspects of *C. elegans* physiology.

## CONCLUSIONS

Here we present a robust and flexible resource for the community that has the potential to bring *C. elegans* research into a more natural and ecologically relevant microbial setting while retaining its strengths as a model system. We demonstrate that the CeMbio strains, either alone or as a community, can affect a key fitness-affecting trait such as developmental rate. Considering that *C. elegans* in nature is inhabited by a diverse microbial community (Dirksen *et al*. 2016; Samuel *et al*. 2016; Berg *et al*. 2016b; Johnke *et al*. 2020), the use of the CeMbio resource in *C. elegans* research will help us to produce a more realistic understanding of nematode biology. We anticipate that the CeMbio community affects the nematode’s interaction with pathogens, as it contains several strains with immune-protective effects, including the two *Pseudomonas* spp. MSPm1 and MYb11, the *Enterobacter* sp. CEent1, and *Ochrobactrum* sp. MYb71 (Montalvo-Katz *et al*. 2013; Berg *et al*. 2016a, 2019; Dirksen *et al*. 2016; Kissoyan *et al*. 2019). Moreover, previous analyses of the *C. elegans* transcriptome response to *Ochrobactrum* sp. MYb71 suggests that the bacteria further affect fertility, energy metabolism, metabolism of specific amino acids, and folate biosynthesis (Yang *et al*. 2019). We expect that other well-studied *C. elegans* phenotypes are also influenced by colonization with these bacteria. By coupling this resource to extensive microbial genomic resources and metabolic models and a small set of bacteria, we anticipate that the CeMbio resource will both provide a facile entry point for *C. elegans* researchers into the more natural world and a nearly limitless arena to explore combinations of these strains together.

## ACKNOWLEDGEMENTS

This work was supported by: NIH grants DP2DK116645 (to B.S.S), R01OD024780 and R01AG061302 (to M.S.); an JGI/DOE grant CSP-503338 (to B.S.S. and H.S.); the Max-Planck Society (fellowship to H.S.); and the German Science Foundation via the CRC 1182 (project A1 to C.K. and H.S.), the Excellence Cluster Inflammation at Interfaces (EXC 306; C.K., H.S.), and under Germany’s Excellence Strategy EXC 2167–39088401 (C.K. and H.S.). Some strains were provided by the CGC, which is funded by NIH Office of Research Infrastructure Programs (P40 OD010440). We also thank Sabrina Butze, Simon Graspeuntner, Georgios Marinos, Katharina Ratjen, Ateequr Rehman, Daniela Vidal and Wentao Yang for support and advice, plus members of the Samuel and Schulenburg labs for helpful comments.

## SUPPLEMENTAL MATERIALS

**Figure S1.**
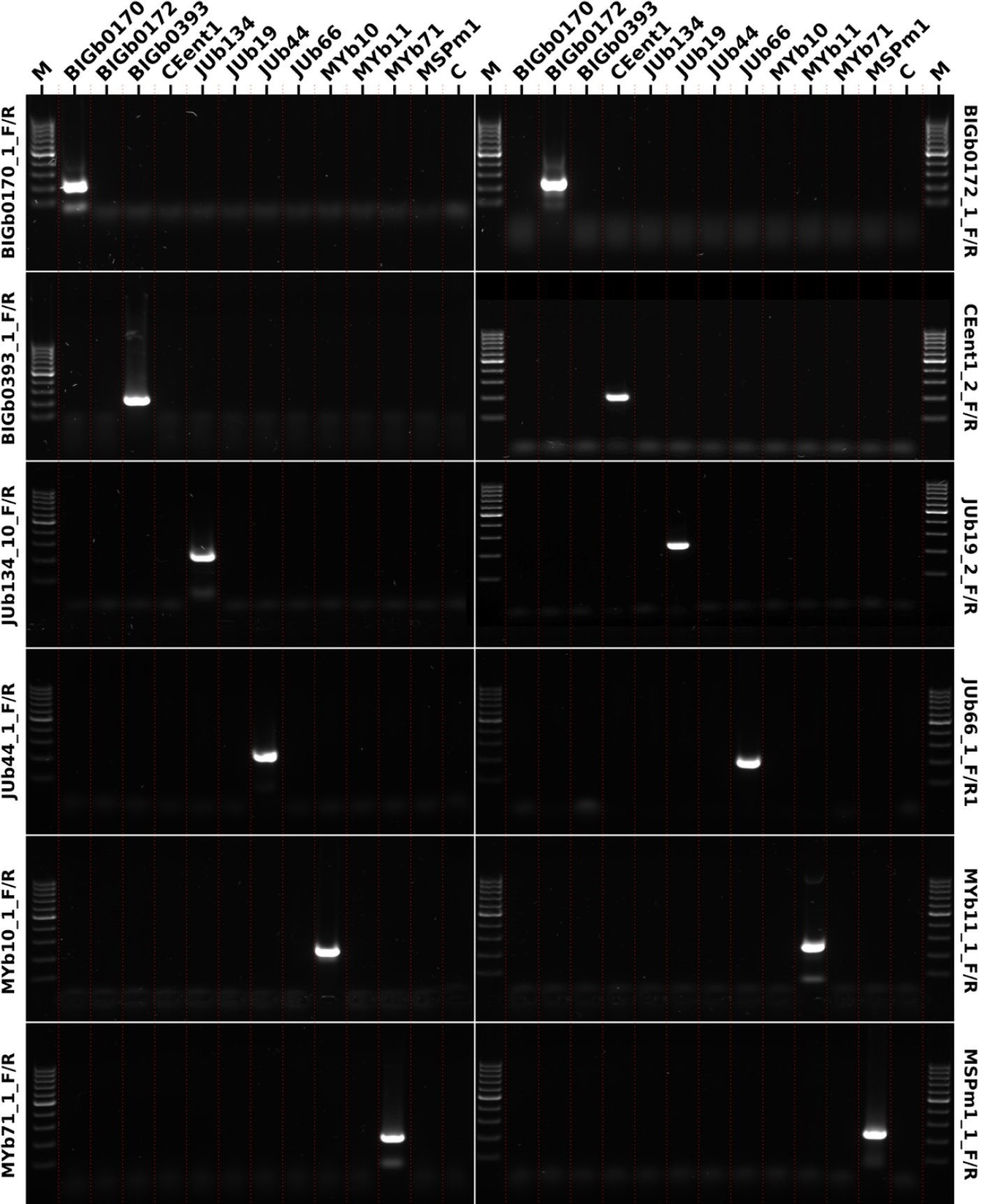
Specificity of the diagnostic PCR primers for the CeMbio strains. To demonstrate the specificity of our diagnostic primers, we conducted PCRs for each primer pair using DNA isolated from pure bacterial cultures of all CeMBio strains. A primer pair was deemed specific if using it resulted in a band of the expected size for only the DNA from the respective target strain, but not for DNA of the other strains. The shown image is a montage of 12 agarose gel electrophoresis pictures (single gels separated by white lines). The lanes contain either a 100 bp ladder (M), water (C), or the DNA of the respective CeMBio strains. The diagnostic primer used in all reactions of a single gel is noted on the side of the image. The images have been scaled to align lanes containing the same DNA and red lines have been added for visual guidance.

**Figure S2.**
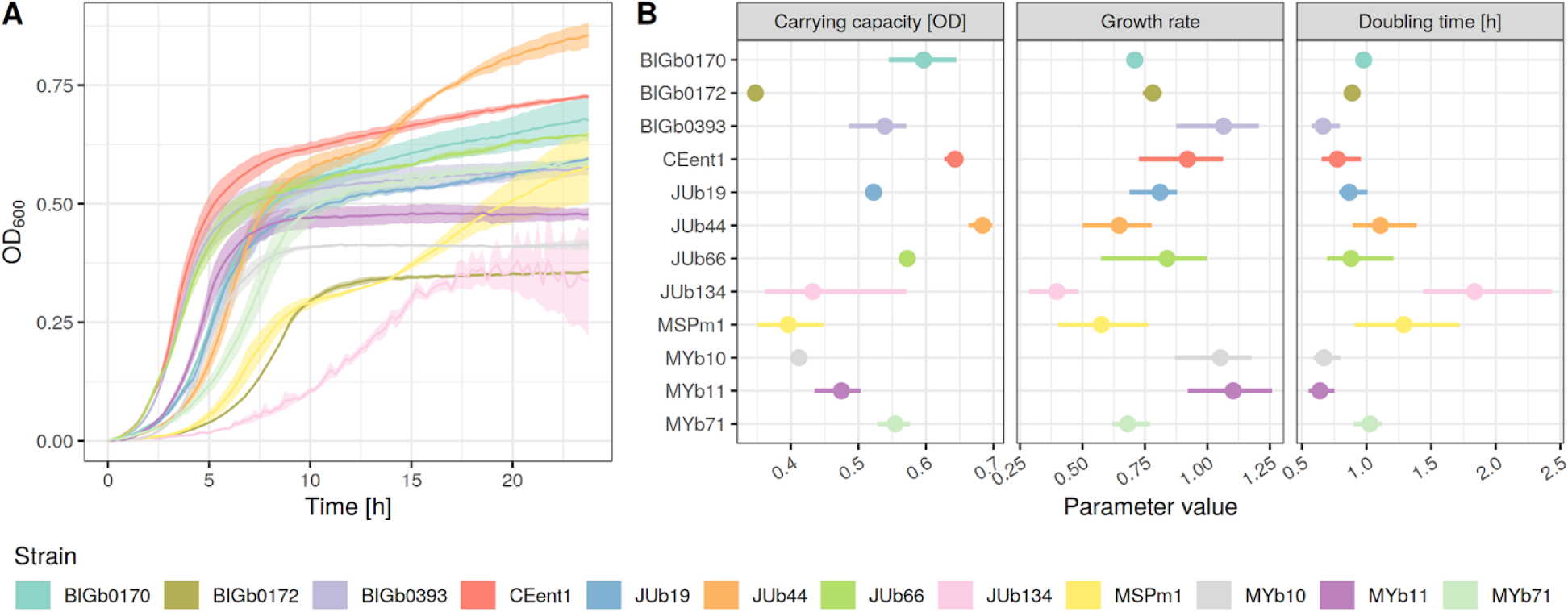
Growth dynamics of the CeMbio strains. **(A)** Growth curves. The bacteria were grown in liquid LB medium at 25°C under aerobic conditions and growth was measured as an increase in optical density at 600 nm for 24 h. **(B)** Carrying capacity, growth rate and doubling time as calculated from (A) using the R package growthcurver. Values are given as median and range [n = 3 replicates / condition].

**Figure S3.**
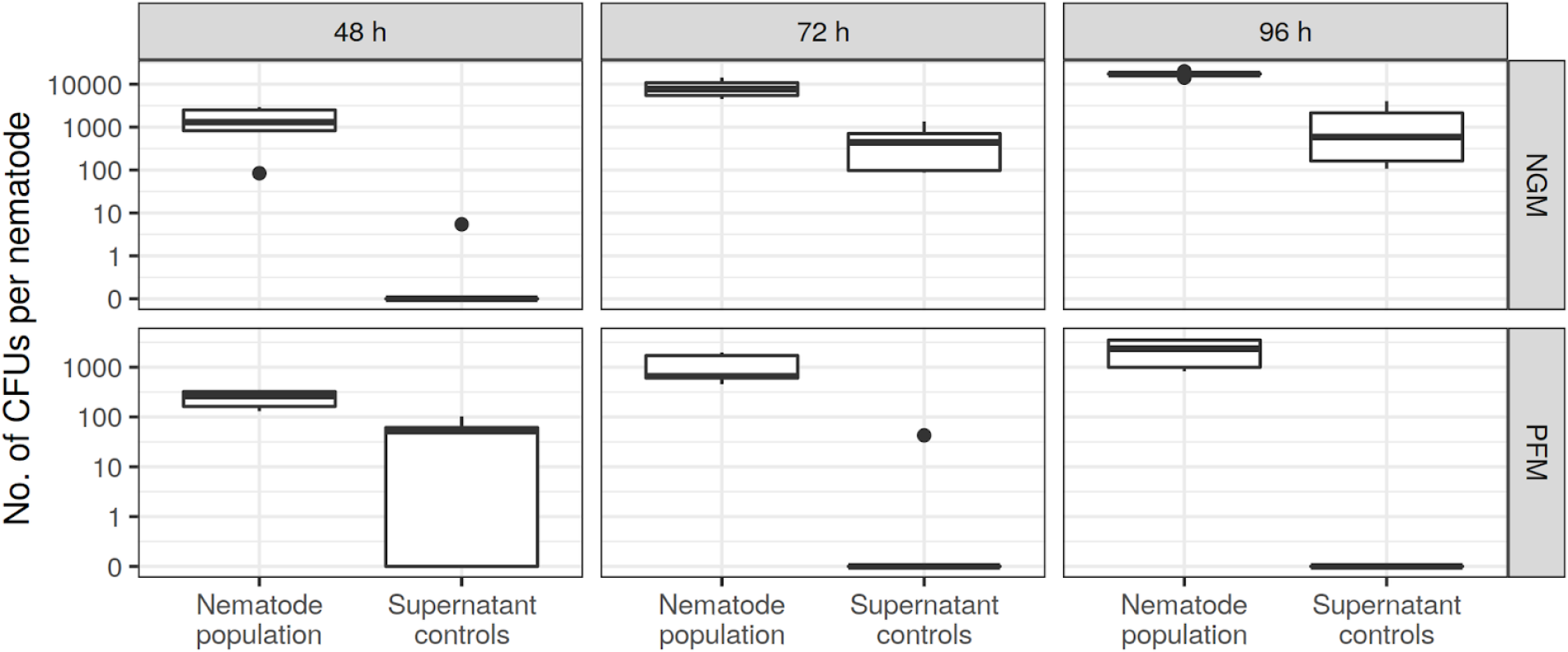
Efficiency of the high-throughput nematode surface sterilization method. We modified our previously established protocol for surface-sterilizing *C. elegans* by combining it with high-throughput separation of nematodes from bacteria (Dirksen *et al*. 2016; Papkou *et al*. 2019). Based on a comparison of the CFUs found in nematodes and the supernatant after the washing procedure, we estimate the carry-over of bacteria from lawns to nematodes during harvest to be ∼5.5%. We repeatedly observed a clear separation of microbiomes from nematodes and plate lawns in a subsequent 16S amplicon analysis (e.g., Figure 5E), indicating that the residual microbes do not mask the nematode microbiome signatures. At the same time, this protocol is significantly faster, allowing the processing of ∼48 samples per hour, and using bleach is bactericidal indiscriminately to potential antibiotic resistances present in the CeMbio bacteria or other bacteria of interest.

**Table S1. List of isolated bacteria from the JUb expanded collection**. This table contains metadata related to the addition of bacteria labelled from JUb130 to JUb274. The table includes the closest relative based on BLAST analysis, taxonomy, isolation origin and 16S rRNA sequences.

**Table S2. A complete list of OTUs from the meta analysis of wild *C. elegans***. The OTU table was generated by clustering 16S rRNA sequences in the V3-V4 region of the 23 wild worm samples (Dirksen et al 2016) at 94% identity. The first column shows the IDs for each OTU identified from the dataset. Column 2 to 24 list raw counts of sequencing reads for each OTU ID in the corresponding 23 *C. elegans* samples. Column 25 to 29 provide taxonomy classification from phylum to genus level for each OTU ID. The last column summarizes commonality of each OTU across all 23 *C. elegans* samples. The 12 OTUs that are 100% common in all worm samples are highlighted in bold.

**Table S3. Diagnostic PCR primers for the CeMbio strains**. All primers are designed to specifically amplify only a single CeMbio strain (Figure S2). Forward primer, reverse primer: Sequence of the forward and reverse primers in 5’ - 3’ direction. Tm: Melting temperature. Size: Size of the amplicon in base pairs. Target gene: Prokka annotation of the gene targeted by the primers.

**File S1. 16S based phylogenies of the CeMbio strains and bacterial type strains belonging to the same family**. The provided archive file contains the 9 tree files generated by *IQ-TREE* (one per bacterial family, see method section on strain characterization for details), as well as a rendering of all trees in a single PDF file, that was used to assign phylogenetic identities to the CeMbio strains.

**File S2. Additional statistics and experimental raw data**. The provided file contains the following tables: OD600 measurements for the CeMbio strain growth dynamics (Figure S2); developmental timing counts (Figure 2); single bacteria colonization data (Figure 5); experiment 3: DESeq2 statistics; experiment 2 - ASV abundances (Figure 3A); experiment 2 - CFU data (Figure 3F); experiment 2 - DESeq2 statistics; experiment 2 - OD600 calibration curves for CFU inference; experiment 2 - CFU inference calculations; experiment 3 - ASV abundances (Figure 4A); experiment 3 - CFU data (Figure 4F and Figure S3); any scripts used for analysis or figure generation.

**File S3. Phylogenomic reconstructions of the CeMbio strains**. The provided file contains additional details and code used to generate the CeMbio phylogenomic reconstruction, as well as partial and complete phylogenomic trees.

**File S4. Revised metabolic models of CeMbio strains**. Adapted genome-scale metabolic models for the 12 CeMBio strains in SBML format. The models available in File S2 were updated according to phenotypic data available from BIOLOG experiments using gapseq.

